# Cadherin Adhesion Complexes Direct Cell Aggregation in the Epithelial Transition of Wnt-Induced Nephron Progenitor Cells

**DOI:** 10.1101/2023.08.27.555021

**Authors:** Balint Der, Helena Bugacov, Bohdana-Myroslava Briantseva, Andrew P. McMahon

**Affiliations:** Department of Stem Cell Biology and Regenerative Medicine, Eli and Edythe Broad Center for Regenerative Medicine and Stem Cell Research, Keck School of Medicine, University of Southern California, Los Angeles, 90033, USA; Department of Urology, Faculty of Medicine, Semmelweis University, Budapest, 1082, HU; Institute of Translational Medicine, Faculty of Medicine, Semmelweis University, Budapest, 1094, HU; Icahn School of Medicine at Mount Sinai, New York, NY 10029, USA

**Keywords:** kidney, nephron progenitor, renal vesicle, Wnt, β-catenin, induction, cadherin cell adhesion

## Abstract

In the developing mammalian kidney, nephron formation is initiated by a subset of nephron progenitor cells (NPCs). Wnt input activates a β-catenin (*Ctnnb1*)-driven, transcriptional nephrogenic program. In conjunction, induced mesenchymal NPCs transition through a pre-tubular aggregate to an epithelial renal vesicle, the precursor for each nephron. How this critical mesenchymal-to-epithelial transition (MET) is regulated is unclear. In an *in vitro* mouse NPC culture model, activation of the Wnt pathway results in the aggregation of induced NPCs into closely-packed, cell clusters. Genetic removal of β-catenin resulted in a failure of both Wnt pathway-directed transcriptional activation and the formation of aggregated cell clusters. Modulating extracellular Ca^2+^ levels showed cell-cell contacts were Ca^2+^-dependent, suggesting a role for cadherin (Cdh)-directed cell adhesion. Molecular analysis identified *Cdh2*, *Cdh4* and *Cdh11* in uninduced NPCs and the up-regulation of *Cdh3* and *Cdh4* accompanying the Wnt pathway-induced MET. Genetic removal of all four cadherins, and independent removal of α-catenin, which couples Cdh-β-catenin membrane complexes to the actin cytoskeleton, abolished cell aggregation in response to Wnt pathway activation. However, the β-catenin driven inductive transcriptional program was unaltered. Together with the accompanying paper (Bugacov *et al*., submitted), these data demonstrate that distinct cellular activities of β-catenin - transcriptional regulation and cell adhesion - combine in the mammalian kidney programs generating differentiated epithelial nephron precursors from mesenchymal nephron progenitors.

**Summary statement:** Our study highlights the role of Wnt–β-catenin pathway regulation of cadherin-mediated cell adhesion in the mesenchymal to epithelial transition of induced nephron progenitor cells.

## Introduction

The developmental morphogenesis of complex tissues requires the coordinated action of distinct cell behaviors (Gumbiner, 1996). In the mammalian kidney, the induction of mesenchymal nephron progenitor cells (NPCs), which initiates a nephrogenic developmental program, is coupled to the establishment of an epithelial nephron precursor, the renal vesicle (RV; (McMahon, 2016). The developmental routines of NPC induction and mesenchymal-to-epithelial transition (MET) continue for days (mouse) to weeks (human) in conjunction with the expansion of the starting pool of NPCs. Consequently, the formation of a species-appropriate complement of nephrons - around 14,000 nephrons in the mouse and one million in the human kidney (McMahon, 2016; Schnell et al., 2022; Short et al., 2014) – is dependent on an orchestrated set of cellular processes. Further, NPC development is closely linked to parallel development of the adjacent ureteric progenitor cells (UPCs) of the collecting system (Shakya et al., 2005), and interstitial (stromal) progenitor cells (Wilson and Little, 2021).

Multiple evidence has highlighted the critical role of canonical Wnt9b/β-catenin (*Ctnnb1*) mediated signaling in the primary induction of NPCs (Guo et al., 2021; Park et al., 2007; Ramalingam et al., 2018). Transduction of UPC-derived Wnt9b signals by a subset of overlying NPC’s results in the accumulation of β-catenin (*Ctnnb1*), which associates with Lef/Tcf DNA-binding complexes, to transcriptionally activate Wnt-targets (Carroll et al., 2005; Park et al., 2012). *In vivo* analysis and *in vitro* modeling of these events in primary NPC culture have identified Lef/Tcf/β-catenin dependent regulators of the nephrogenic program and demonstrated the direct interaction of Lef/Tcf/β-catenin complexes with cis-regulatory modules regulating target gene expression (Guo et al., 2021; Park et al., 2012). In parallel with transcriptional activation of the nephrogenic program, induced NPCs cluster and condense into pre-tubular aggregates (PTAs), which complete a MET establishing epithelial renal vesicles, precursors to the functional nephrons of the mammalian kidney (Georgas et al., 2009).

The molecular mechanism governing morphological transition of mesenchymal NPCs into an epithelial nephron anlagen are unclear. β-catenin plays a critical role in cadherin-mediated cell adhesion complexes (Halbleib and Nelson, 2006) and is consequently an attractive candidate for promoting the aggregation and epithelial transition of NPCs. However, constitutive elevation of β-catenin levels blocks epithelial formation *in vitro* (Park et al., 2012) and several lines of evidence suggest a non-canonical signaling role for Wnt4, a primary transcriptional target of Wnt9b/b-catenin induced NPCs in the MET (Saulnier et al., 2002; Stark et al., 1994; Tanigawa et al., 2011; Vainio et al., 1999). Further, in the gastrulating vertebrate embryo (Kelly et al., 2004; Liu et al., 1999) and metastasis of a variety of epithelial cancers (Dongre and Weinberg, 2019; Huang et al., 2022) canonical Wnt signaling is linked to an opposite cellular program: an epithelial-to-mesenchymal transition (EMT).

Here, we used cell imaging, cell profiling and molecular-genetic approaches to characterize the morphological changes associated with Wnt-β-catenin mediated induction of NPCs *in vitro*. These studies, together with those in the accompanying paper (Bugacov *et al*., submitted), demonstrate that distinct actions of β-catenin in transcriptional regulation and cell-cell adhesion coordinate gene regulatory and morphological cellular programs directing early mammalian nephrogenesis.

## Results

### Increasing Wnt/β-catenin activity in NPCs in vitro models NPC induction and morphogenesis in vivo

To examine the mechanisms of canonical Wnt pathway action on NPC programs, we used an *in vitro* model comprising highly purified naïve uninduced NPCs cultured in a defined Nephron Progenitor Expansion Media (NPEM). NPEM replicates multiple signaling activities linked to the maintenance and expansion of NPCs in the mouse kidney (Brown et al., 2015). As shown in the accompanying manuscript (Bugacov *et al*., submitted), NPC fate outcomes in this model are dependent on the concentration of CHIR99021 (hereafter CHIR), a small molecule antagonist of the serine-threonine kinase, GSK3β (**Fig. 1A–B**). As, GSK3β-mediated phosphorylation of β-catenin results in β-catenin turnover by a destruction complex (Bennett et al., 2002), varying degrees of stabilization of β-catenin mirror dose-dependent, Wnt-receptor-mediated canonical Wnt signaling (**Fig. 1B**). Low CHIR (1.25 µM) stimulated the proliferation of dispersed NPCs which maintained a Six2^high^/Jag1^−^ NPC state (**Fig. 1C–D**). While high CHIR (5 µM) induced the transcriptional activation of a nephrogenic program (see Bugacov *et al*., submitted for more complete analysis) and the aggregation of induced Six2^low^/Jag1^+^ NPCs into tightly adherent multicellular clusters (**Fig. 1C–D**). Aggregates showed enhanced cell membrane associated accumulation of β-catenin in a continuous ring **(Fig. 1E**; arrowed in **Fig. 1H-–I**). Consequently, nearest cell neighbour distance decreased in high CHIR (**Fig. 1F**), in conjunction with induction of Jag1, an early and direct transcriptional response to canonical Wnt complexes (**Fig. 1 G;** (Guo et al., 2021). While nuclear volume was unaltered in high CHIR induced aggregates (**Fig. 1J**), nuclear height and cell height increased (**Fig. 1K–L**) and overall cell volume decreased (**Fig. 1M**). Thus, 24 h following a switch from low to high CHIR, NPCs morphologically transform from flattened mesenchyme to columnar, multicellular aggregates, reflective of PTA formation *in vivo*.

**Figure 1.**
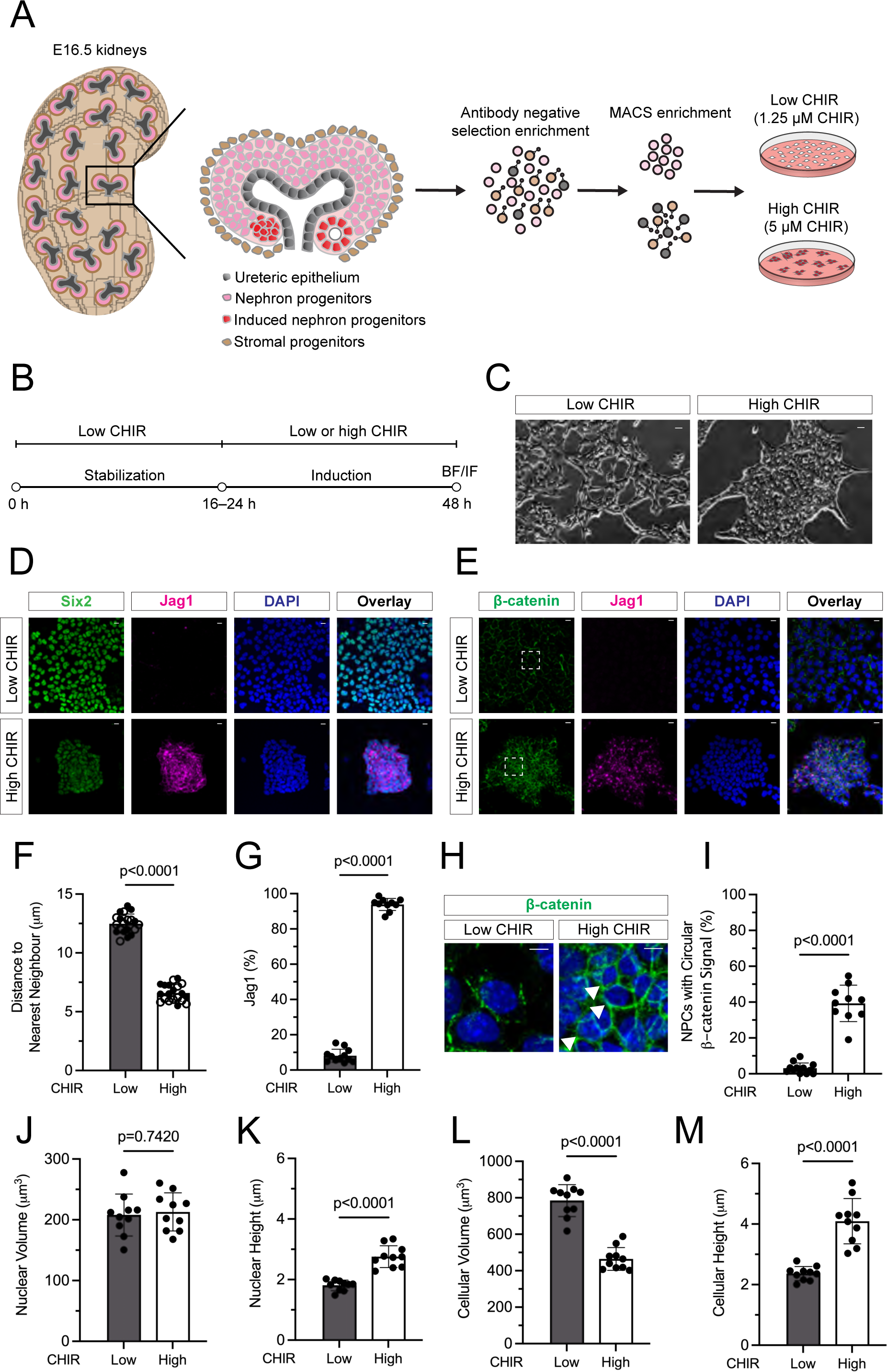
NPCs undergo morphological and transcriptional changes in vitro as a result of increased Wnt stimulus. (A) Schematic representation of NPC isolation and culture system in NPEM supplemented with low (1.25 μM) CHIR or high (5 μM) CHIR. (B) Schematic diagram of experimental protocol for panels in Fig.1. BF: brightfield, IF: immunofluorescence (C) Phase contrast images of dispersed NPCs in low CHIR condition vs. NPC aggregates in high CHIR condition. Scale bars are 50 μm. (D) IF staining of isolated E16.5 NPCs characterizing Six2 (green), Jag1 (magenta) and nuclear DNA (DAPI-blue). Scale bars are 10 μm. (E) IF staining of isolated E16.5 NPCs characterizing β-catenin (green), Jag1 (magenta) and DAPI (blue). Squared area is magnified in Fig. 1H. Scale bars are 10 μm. (F) Quantification of cell aggregation by plotting distance to nearest neighbor. Unpaired *t* test. 2 technical replicates plotted as differently filled circles. (G) Quantification of induction by the percentage of Jag1^+^ immuno-positive NPCs. Unpaired *t* test. (H) Representative images of continuous membrane Ctnnb1 (green, arrowheads) in aggregated NPCs in high CHIR condition (DAPI^+^ nuclei, blue). In low CHIR condition, Ctnnb1 is discontinuous at the cell membrane. The areas of the high magnification images are highlighted on Fig. 1E. (I) Quantification of Ctnnb1 membrane distribution. Unpaired *t* test. (J–M) Quantification of cell morphology changes of E16.5 WT NPCs cultured in low CHIR and high CHIR conditions. Statistical tests: nuclear volume (unpaired *t* test); nuclear height (unpaired *t* test); cellular volume (unpaired *t* test); cellular height (unpaired *t* test).

### High resolution timelapse imaging of cell contact stabilization on induction of NPCs

To investigate the dynamics of cell aggregation, NPCs labeled with a cell membrane targeted tdTomato fluorescent protein (Muzumdar et al., 2007) were cultured in the absence of CHIR, or in low or high CHIR conditions and followed by time lapse confocal microscopy over 6h. Increasing CHIR levels resulted in increased clustering of NPCs: clustering was tightest and cell-cell adhesions most stable in high CHIR conditions (**Vid. S1, Fig. S1A**). Consequently, NPCs cultured in high CHIR showed the shortest average distance to nearest neighbors **(Fig. S1B)** and experienced fewer contacts per cell over the imaging time course **(Fig. S1C)**. Thus, NPCs are more motile and maintain shorter cell-cell contacts in both the absence of CHIR or in low CHIR, while high CHIR conditions stabilized cell-cell contacts (**Fig. S1D**). In all conditions, NPCs send out filopodia resulting in cell-cell contacts. An increased membrane surface contact in high CHIR (**Fig. S1E**) correlated with an increased stability of filopodia interactions between cells in high CHIR which was evident as early as 3h after initiation of high CHIR culture (**Fig. S1. F–H**).

### β-catenin activity is required for cell-cell aggregation of induced NPCs

To examine the requirement for β-catenin in cell-cell aggregation, RNA-lipofection was used to remove *Ctnnb1* function and β-catenin activity by either sgRNA/Cas9-directed gene knockout (KO) or CRE-directed critical exon excision, identifying genetically modified cells through co-activation of an mCherry reporter (**Fig. 2A**; for details see accompanying manuscript Bugacov *et al*., submitted). As expected, loss of β-catenin blocked high CHIR mediated induction of Jag1, but also resulted in a failure in cell-cell aggregation (**Fig. 2B, C–F**). Exclusion of mutant cells could be observed in 6–12 h post addition of high CHIR (**Fig. S2, A–B,** live imaging in **Vid. S2**). Addition of a bi-specific antibody (BSAB), which mediated direct activation of cell surface Wnt-receptors (Janda et al., 2017), gave an indistinguishable outcome to high CHIR (**Fig. S2B)**. In contrast to β-catenin KO, when NPC induction was blocked by the KO of all four Tcf/Lef factors mediating β-catenin’s transcriptional activity, uninduced NPCs (Jag1^−^) were retained in aggregates alongside induced wild-type NPCs (Jag1^+^; **Fig. 2G).**

**Figure 2.**
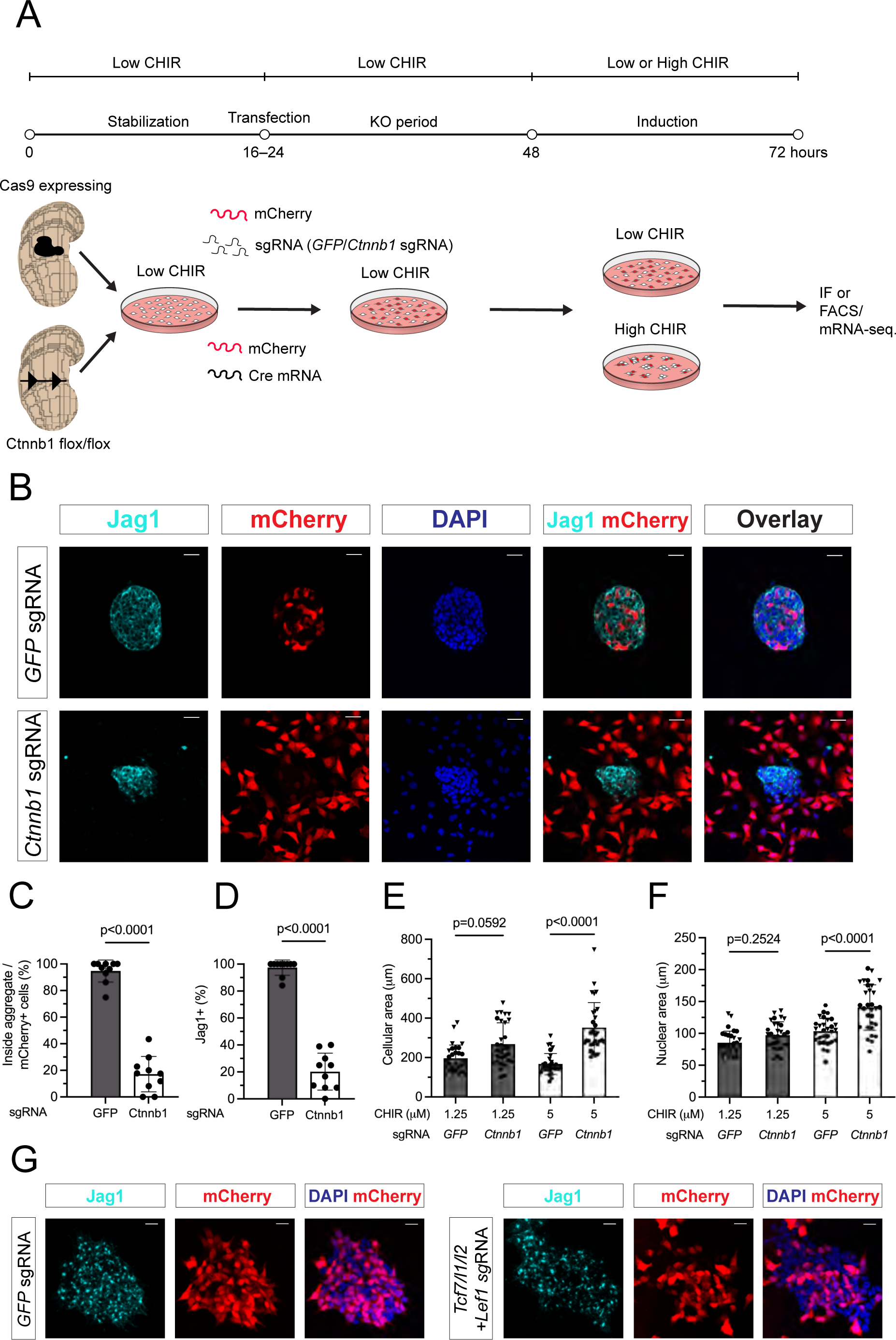
In vitro deletion of ß-catenin results in NPC cell sorting and abolished the CHIR dependent induction program. (A) Schematic of the experimental protocol in β-catenin KO experiments with 24 h KO period and 24 h induction time for Fig. 2B–F. (B) Representative images of CTRL (GFP sgRNA) and β-catenin-KO (Ctnnb1 sgRNA) conditions showing loss of cell aggregation and Jag1 induction in mCherry^+^ NPCs on loss of Ctnnb1 in high CHIR conditions. Scalebars are 25 μm. (C, D) Quantification of cells sorting phenomenon by % of cells within aggregates and induction by percentage of Jag1^+^ NPCs as a result of knocking out β-catenin in NPCs. Statistics: Mann-Whitney test. (E, F) Quantification of changes in cellular morphology as a result of β-catenin-KO in NPCs. Statistics: cellular area (ordinary one-way ANOVA), nuclear area (Kruskall-Wallis test). (G) Representative images of CTRL (GFP sgRNA) and Tcf/Lef-KO (Tcf7/Tcf7l1/Tcf7l2/Lef1 sgRNA) conditions after 48 h KO period showing transfected cell distribution and alterations in induction (Jag1). Scalebars are 20 μm.

To obtain a better insight into the dynamics of β-catenin-KO NPCs, we focused on control gRNA and β-catenin gRNA transfected, β-catenin-KO NPCs at the edges of colonies 6-12 h into high CHIR induction (**Vid. S3, Fig. S2C**). While wild-type NPCs in control transfections maintained adherence to the larger cell aggregate, β-catenin-KO NPCs did not form longer term stable contacts, and moved along the edges of cell aggregates, eventually moving away from cell-cell aggregates. In summary, β-catenin activity was required for stabilizing cell-cell associations that were essential for the generation of multicellular aggregates in conjunction with transcriptional activation of canonical Wnt signaling.

### Functional analysis of α-catenin supports independent actions of β-catenin in transcriptional regulation and aggregation of NPCs

In Ca^2+^ mediated, cadherin directed cell-cell adhesion, β-catenin interactions with membrane bound cadherins are relayed to the cytoskeleton through *α*-catenin (Aberle et al., 1994; Maiden and Hardin, 2011). Culturing NPCs in media without Ca^2+^, in either low or high CHIR conditions, led to a rapid loss of cell-cell contacts within 5 minutes (**Fig. S2 D–E**) which could be reversed by the addition of Ca^2+^, though over a longer time course (30 minutes). The results support a role for Ca^2+^-dependent cadherin-catenin complex in aggregation of NPCs though it should be noted that Ca^2+^ removal has been shown to also influence integrin functions (Kirchhofer et al., 1991).

Next, we examined a potential role for *α*-catenin in the MET. *In vivo* in developing E16.5 kidney, β-catenin and α-catenin co-localize within Jag1^+^ cells in induced aggregating NPCs (**Fig. S3A**). To examine the function of α-catenin, we used α-catenin sgRNA and gene editing in Cas9+ NPCs to remove the function of the α-catenin encoding gene, *Ctnna1*, extending the KO period to 48 h for efficient removal of the α-catenin protein (∼95% removal; **Fig. S3C-D**). Strikingly, while α-catenin removal blocked cell aggregation in high CHIR, similar to the removal of β-catenin, α-catenin KO cells induced Jag1, indicative of an active canonical Wnt-directed transcriptional response (**Fig. 3D–G**). Thus, α-catenin separates distinct actions for β-catenin in the NPC inductive program. Replacing high CHIR with Wnt3a gave a similar inductive response (**Fig. S3E**) and cell distribution indicating that the α-catenin dependent cell aggregation was a *bona fide* outcome of canonical Wnt pathway activation (**Fig. 3H–I**).

**Figure 3.**
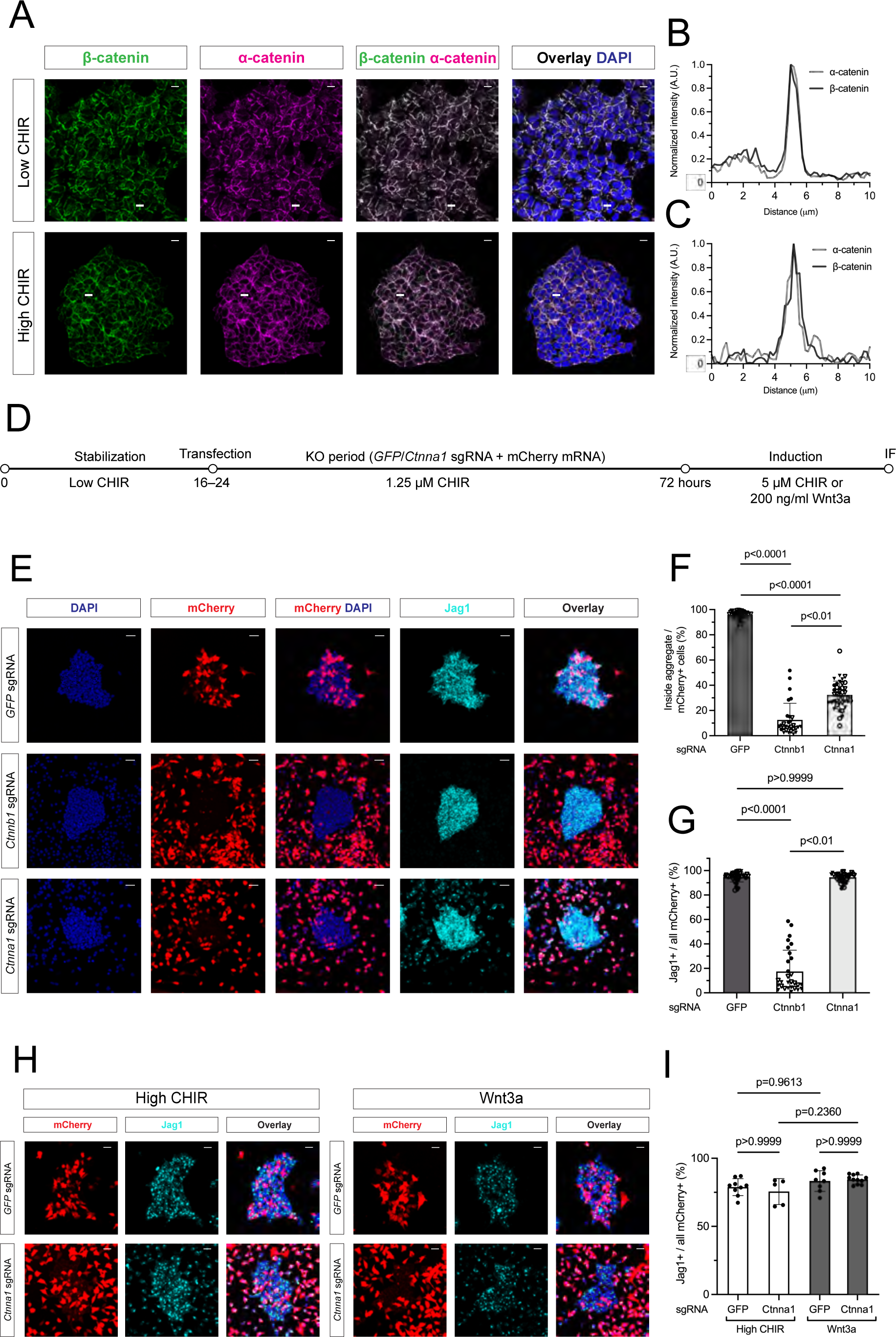
Removal of α-catenin in NPCs in high CHIR conditions phenocopies β-catenin KO-dependent cell sorting but induction of Jag1 was not altered. (A) Representative images of isolated E16.5 NPCs IF co-staining of α- and β-catenin (α-catenin: magenta, β-catenin: green) in low and high CHIR conditions. Intensity values over the in-frame lines are plotted on B–C. Scale bars are 10 μm. (B–C) Normalized fluorescent intensity (normalization to baseline values) showing highly overlapping intensity values for α- and ß-catenin. (D) Schematic of the experimental protocol in α-catenin KO experiments corresponding to Fig. 2E–I. (E) Representative images of CTRL (GFP sgRNA), ß-and α-catenin-KO conditions showing cellular behavior (mCherry, DAPI) and inductive response (Jag1) in 5 μM CHIR. Scalebars are 25 μm. (F, G) Quantification of percentage of Jag1^+^ cells in aggregates in high CHIR analyzed by Kruskal-Wallis test with biological replicates represented by different symbol shapes and technical replicates by different color filling of symbols). (H) Representative images of CTRL (GFP sgRNA) and α-catenin-KO conditions showing altered cell aggregation but normal Jag1 (cyan) induction in α-catenin-KO transfected (mCherry) cells (DAPI: blue nuclear staining) treated with 200 ng/ml Wnt3a. Scalebars 20 μm. (I) Quantification of inductive response in Fig. 3H: percentage of transfected mCherry^+^/Jag1^+^ NPCs. Kruskall-Wallis test.

### Multiple cadherins mediate NPC aggregation on canonical Wnt pathway activation

Together the findings above suggest a role for cadherin-catenin complexes in the cell-cell aggregation of induced NPCs. Previous reports examining single and multiple allelic combinations of cadherin mutants (Cdh2^+/−^, Cdh3^−/−^, Cdh4^−/−^, Cdh6^−/−^) failed to identify a compelling role for cadherins in the MET of NPCs *in vivo* (Mah et al., 2000). To characterize expression of cadherins, we performed *in vitro* bulk mRNA-sequencing of NPCs in low and high CHIR. *Cdh2* and *Cdh11* were expressed at high levels in low CHIR, and downregulated on induction in high CHIR (**Fig. 4A**). In contrast, *Cdh3*, *Cdh4* and *Cdh6* were upregulated in high CHIR conditions (**Fig. 4A**). Immunofluorescent (IF) staining of cultured NPCs, confirmed the presence of Cdh2, Cdh4 and Cdh11 at the cell membrane before induction and the appearance of Cdh3 and downregulation of Cdh11 at the membrane, on high CHIR-mediated induction of NPCs (**Fig. 4B–E**). In contrast, Cdh13 was not detected and Cdh6 was mostly confined to the GM130^+^ Golgi apparatus in induced NPCs, arguing against a role for Cdh6 in cell surface accessible adhesion complexes. (**Fig. S4A–C**). Immuno-analysis confirmed key features of predicted cadherin distribution in the NPC lineage in the developing kidney *in vivo* (tdTomato+ cells in **Fig. 4F–I, Fig. S4D**). A view of cadherin expression in the early human nephrogenic program showed a broadly similar expression for mouse and human cadherin genes (**Fig. 4J, Fig. S4E**).

**Figure 4.**
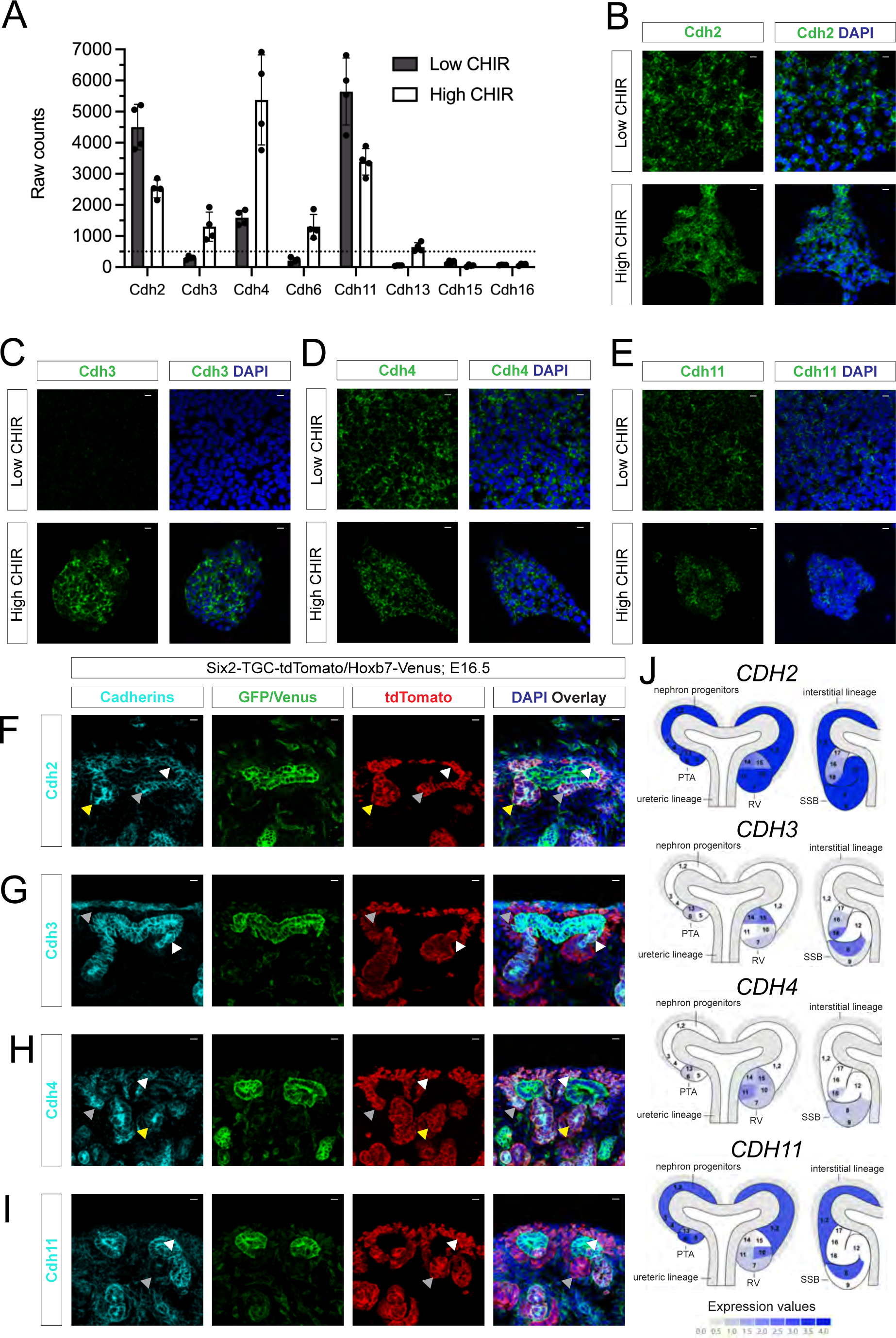
In vitro and in vivo analysis of cadherin mRNA and protein levels reveals that NPC culture system models the in vivo conditions. (A) Raw counts of bulk RNA-seq for selected cadherin mRNAs in E16.5 NPCs following 24 h of culture in low and high CHIR conditions. (B–E) IF staining of isolated E16.5 NPCs showing Cdh2 (B) and Cdh4 (D) in the cell membrane in low and high CHIR conditions, high levels of Cdh3 (C) restricted to the high CHIR condition, and weak membrane Cdh11 labeling (E) in low and high CHIR conditions. Scale bars are 10 μm. (F–I) Representative images of IF staining of E16.5 Six2-TGC-tdTomato/Hoxb7-Venus mouse kidneys highlighting indicated cadherins (cyan), tdTomato (nephron lineage, red) and GFP: nuclear GFP highlights Six2 in NPCs while membrane GFP labels Venus reporter in the ureteric lineage. (G) Cdh2 is present in both uninduced and induced NPCs, and late RVs (white, gray and yellow arrowheads, respectively). (H) Cdh3 is present in the distal segment of late RV including the invading cells of the ureteric bud tip, but not in the uninduced NPCs (white and gray arrowheads, respectively). (I) Cdh4 is present in uninduced NPCs and at higher levels in the PTA and SSB (white, gray and yellow arrowheads, respectively). (J) Cdh11 is present at low levels in uninduced NPCs and levels decrease in the RV (white and gray arrowheads, respectively). Scale bars are 10 μm. (J) Human Nephrogenesis Atlas views (Lindström et al., 2021) of human CDH gene expression in early development of the human nephron lineage. PTA: pre-tubular aggregate, RV: renal vesicle, SSB: S-shaped body

To test a potential role for the cadherins of interest, we identified effective sgRNA KOs for each cadherin demonstrating complete removal of detectable Cdh2 and Cdh11 prior to high CHIR addition 48 h post transduction, and 90-95% of Cdh3 and Cdh4 removal, assaying post induction (**Fig. 5A–J**). Similar efficiencies were observed combining gRNAs for multiple cadherin removals (**Fig. S5A–C**). Whereas targeting of individual cadherins did not alter the aggregation process or inductive process (**Fig. S6A–E**), combinatorial removal of Cdh2, Cdh4 and Cdh11 resulted in an interesting phenotype. Triple KO cells clustered at the boundary of cell aggregates in high CHIR in contrast to control GFP gRNA KO cells, or single cadherin KO cells, which dispersed randomly in the aggregate (**Fig. 6A–C).** Further, membrane levels of Cdh3 were elevated on depletion of Cdh2, Cdh4, Cdh11, consistent with a competitive stabilization of cadherins in the membrane (**Fig. S6F–G**). Strikingly, combining KO of Cdh3 with Cdh2, Cdh4 and Cdh11 (QCKO), prevented cell aggregation in high CHIR **(Fig. 6A–C).** And, similar to α-catenin removal and distinct from β-catenin KO, QCKO cells were Jag1^+^ indicating a β-catenin-dependent transcriptional response in non-aggregated NPCs (**Fig. 6A–D**). Together, these results provide strong evidence for classic cadherin complexes in regulating the MET in the nephrogenic program (**Fig. 6E–F**).

**Figure 5.**
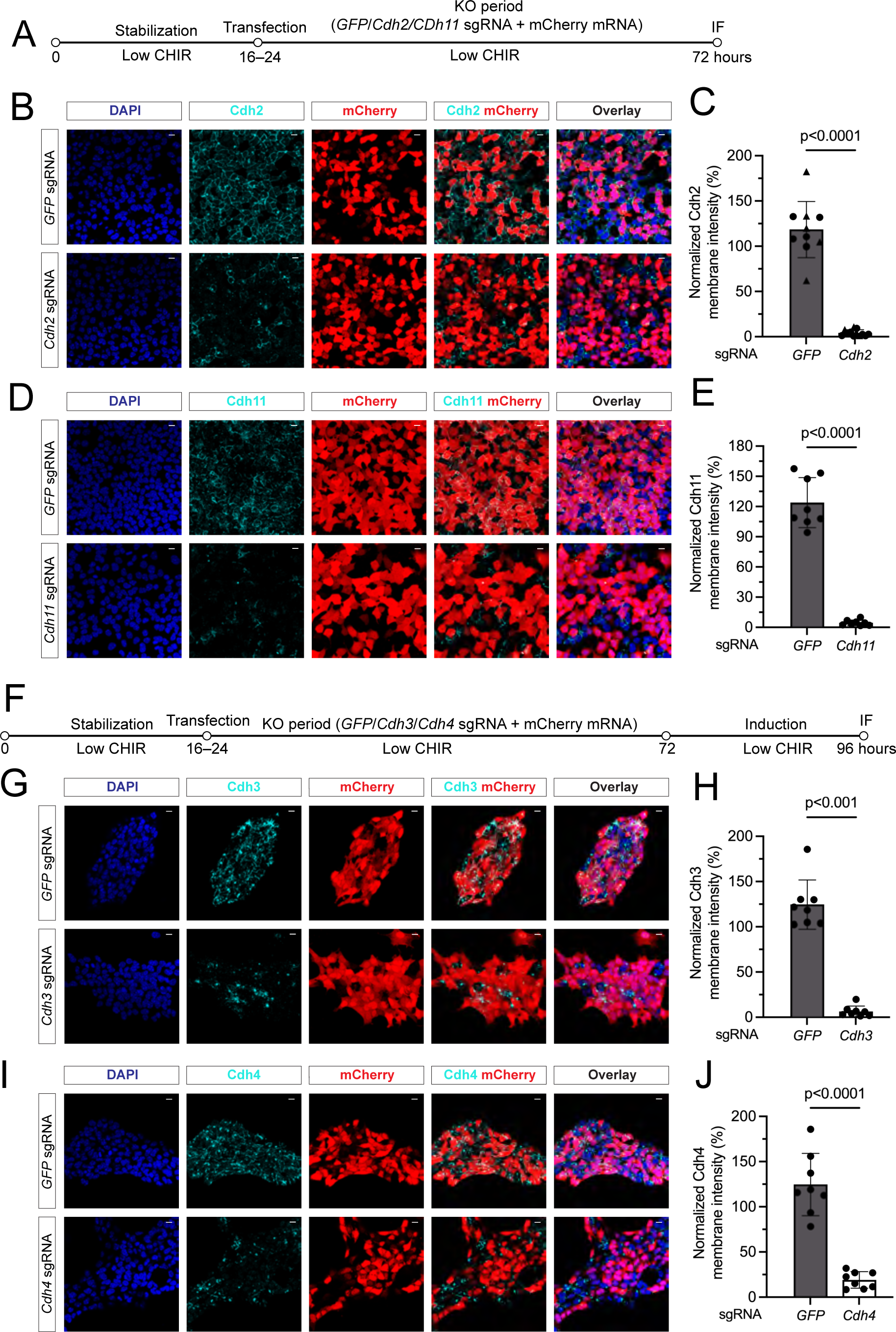
Confirming the removal of individual cadherin proteins by the Cas9-sgRNA KO system. (A) Schematic representation of the experimental protocol to remove Cdh2 and Cdh11 with 48 h KO period. (B) Representative IF images of Cdh2 removal (cyan) in mCherry transfected cells (red) with nuclear staining DAPI (blue). Scale bars are 10 μm. (C) Quantifications of the membrane intensity of Cdh2 KO cells. Unpaired *t* test. (D) Representative IF images of Cdh11 removal (cyan) in mCherry transfected cells (red) with nuclear staining DAPI (blue). Scale bars are 10 μm. (E) Quantifications of the membrane intensity of Cdh11 KO cells. Unpaired *t* test. (F) Schematic representation of the experimental protocol to remove Cdh3 or Cdh4 including 48 h KO period and 24 h induction. (G) Representative IF images of Cdh3 removal (cyan) in mCherry transfected cells (red) with nuclear staining DAPI (blue). Scale bars are 10 μm. (H) Quantifications of the membrane intensity of Cdh3 KO cells. Mann-Whitney test. (I) Representative IF images of Cdh4 removal (cyan) in mCherry transfected cells (red) with nuclear staining DAPI (blue). Scale bars are 10 μm. (J) Quantifications of the membrane intensity of Cdh4 KO cells. Unpaired *t* test.

**Figure 6.**
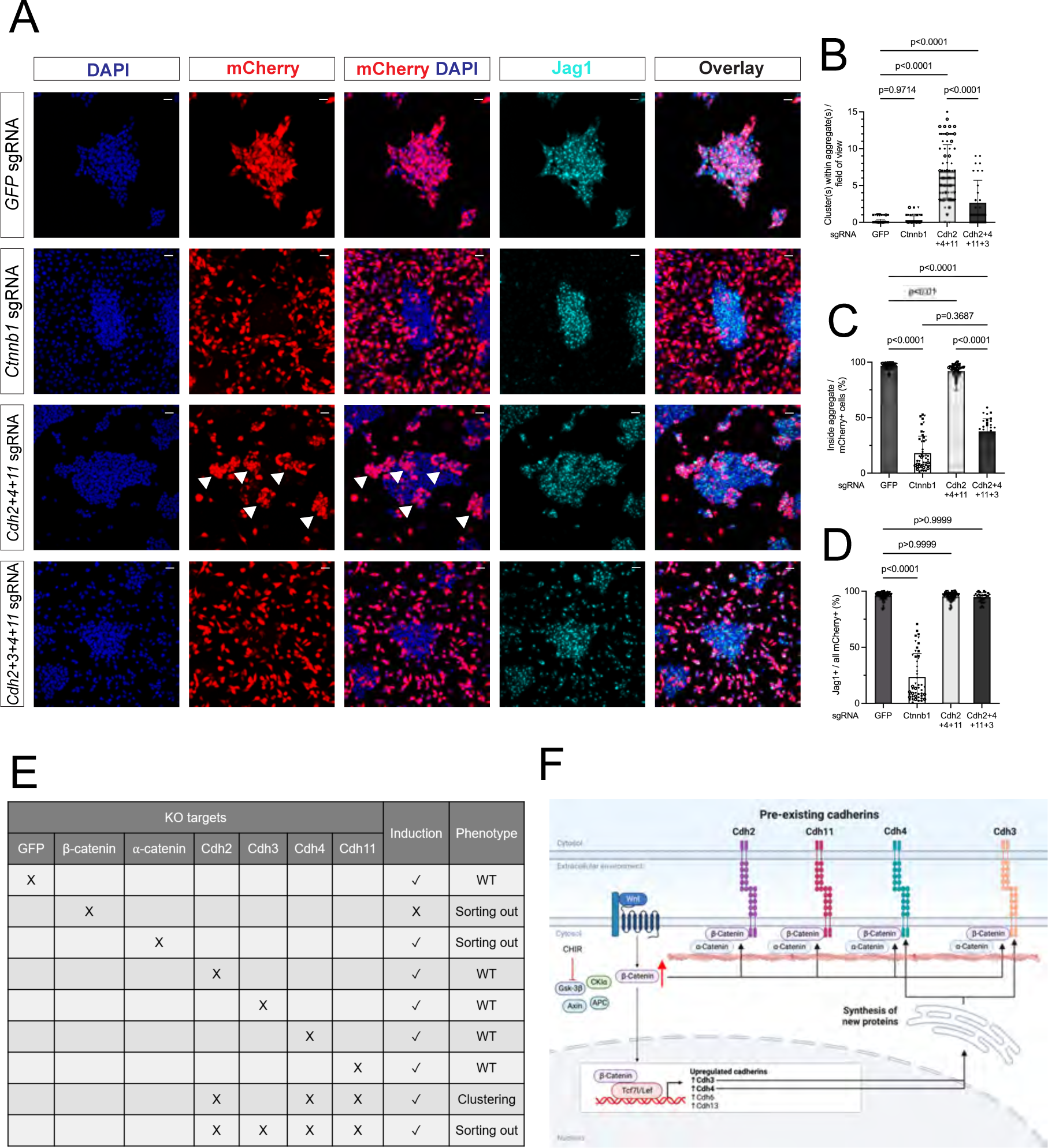
Compound cadherin removal inhibits cell adhesion phenocopying α- and ß-catenin removal, however maintains NPC transcriptional induction. (A) Representative images of CTRL (GFP sgRNA), β-catenin-KO (positive CTRL) and pre-existing cadherin KO (Cdh2, Cdh4, Cdh11), and QCKO (Cdh2, Cdh3, Cdh4, Cdh11) showing cellular behavior (mCherry, DAPI) and induction (Jag1). Arrowheads highlight transfected NPCs clustering within aggregates. Scalebars are 25 μm. (B–D) Quantification of changes in cell adhesion as the number of transfected cell cluster(s) within aggregates (B), cells sorting phenomenon by % of cells within aggregates (C) and induction by Jag1 expression percentage of NPCs in cadherin KO experiments (D). B: ordinary one-way ANOVA, C, D: Kruskall-Wallis test. 3–6 biological replicates (different symbol shapes), 1–2 technical replicates (different fill colors of symbols). (E) Summary table of the effects of single and compounded cadherin-catenin complex KOs on induction and cellular behavior. (F) Proposed mechanism of the NPC aggregation regulated by Wnt/β-catenin via the cadherin-catenin complex modeling the initial step of nephrogenesis *in vivo*.

### Compound cadherin or α-catenin removal does not alter the transcriptional program within induced NPCs

To examine whether induction was independent of cell aggregation, we performed bulk mRNA-sequencing on two biological replicates of FACS sorted genetically modified NPCs utilizing the 48 h KO period followed by 24 h in high CHIR (**Fig. 7A**; **Tables 1.1-4**). Principal component analysis comparing control (CTRL) samples targeting GFP with α-catenin KO, β-catenin KO and QCKO showed similar co-clustering amongst CTRL, α-catenin KO and QCKO experimental samples in each CHIR condition (**Fig. 7B**). In line with findings in the accompanying paper (Bugacov *et al*., submitted), CTRL KO samples targeting GFP showed an expected inductive response with the downregulation of NPC associated genes and upregulation of large set of genes including well recognized Wnt targets such as *Jag1*, *Lef1, Wnt4* and *Lhx1* (Guo et al., 2021; **Fig. 7C**; **Table 1.1**). Comparing induced (high CHIR) CTRL KO and α-catenin KO, only *Ctnna1* was identified as a differentially expressed gene (DEG; absolute log_2_FC cut off 0.5; p-adj cut off 0.5; **Table 1.2)**. In CTRL KO vs. QCKO. the only DEG was *Rasl11a* (**Table 1.2–4**). Thus, cell aggregation did not alter the β-catenin-dependent transcriptional response to high CHIR significantly leading to the conclusion that induction of the nephrogenic program was independent of cell reorganization, in the experimental model.

**Figure 7.**
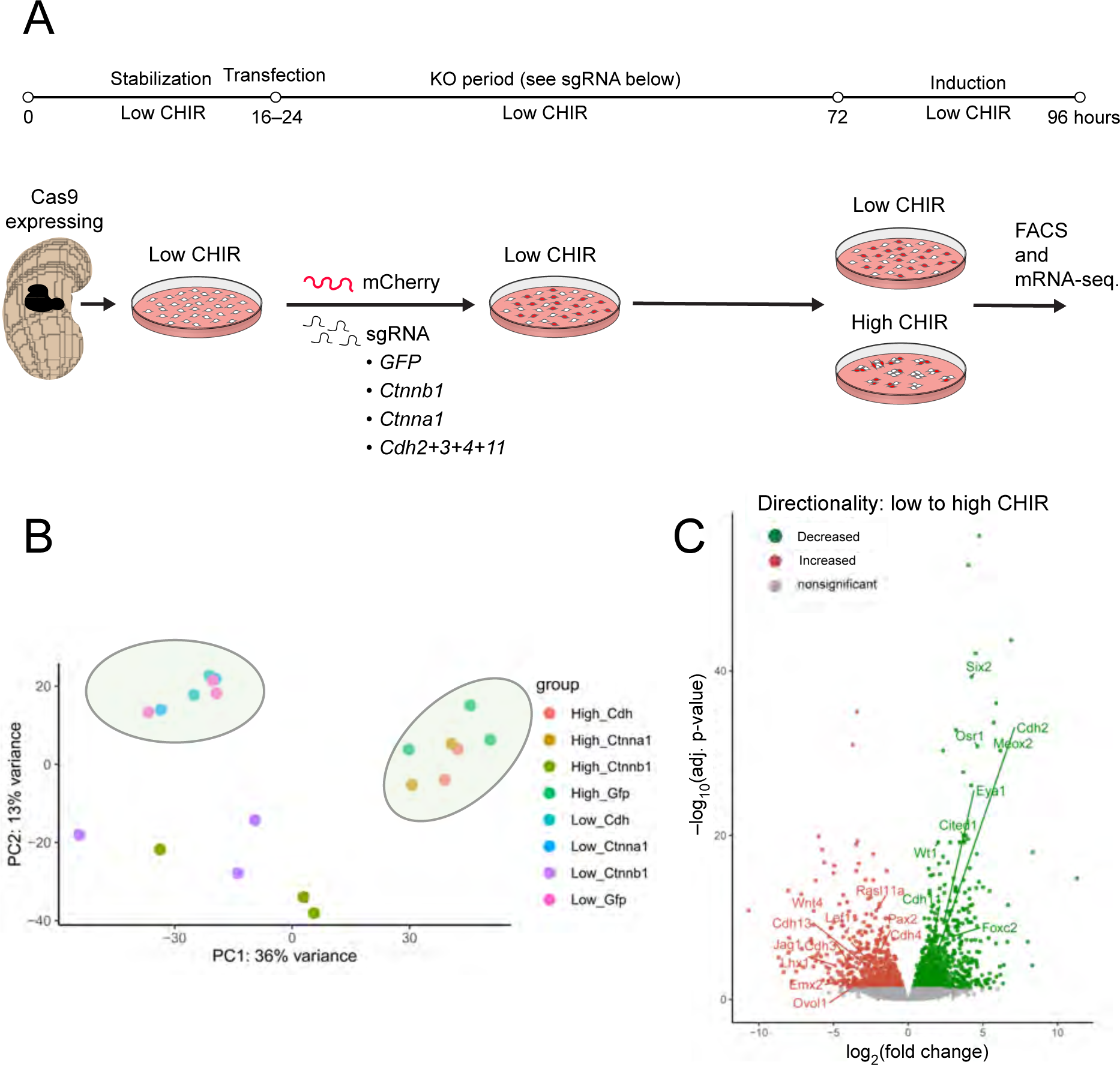
Induction of NPCs is independent of cadherin-mediated cell clustering in NPC culture. (A) Schematic representation of workflow for transcriptional profiling of *Ctnna1, Ctnnb1,* and combined *Cdh2*, *Cdh3*, *Cdh4*, and *Cdh11* knock out (QCKO) in NPC culture. (B) PCA plot of mRNA-seq transcriptional profiles of CTRL GFP KO (GFP sgRNA), β-catenin-KO (*Ctnnb1* sgRNA), α-catenin-KO (*Ctnna1* sgRNA) and QCKO in low and high CHIR conditions. PC1 (36%) variance relates to transcriptional induction status. (C) Volcano plot of low CHIR vs. high CHIR DeSEQ2 bulk RNA-seq analysis examining control GFP gRNA samples in low and high CHIR (corresponding **Table S1.1**). Highlighted genes include self-renewal associated genes-of-interest downregulated in high CHIR (green: *Six2, Cited1, Eya1*) and induction associated genes-of-interest up-regulated in high CHIR (red: *Wnt4, Jag1, Lef1, Cdh3, Cdh4, Lhx1, Ovol1, Emx2)*.

## Discussion

Employing a culture system that reproduces key cellular behaviors and gene regulatory controls associated with Wnt-directed initiation of mammalian nephrogenesis, we provide multiple lines of evidence supporting a critical role for cadherin adhesion complexes in the MET underpinning morphogenesis and patterning of the nephron. Cell-cell contacts between NPCs were Ca^2+^ sensitive and stabilized in high CHIR in a β-catenin process requiring α-catenin and the partially redundant activities of four cadherins: Cdh2, 3, 4 and 11. Examining cell-cell interactions in low and high CHIR suggests uninduced and induced NPCs are actively send out filopodia making new cell contacts, consistent with *in vivo* observations (Combes et al., 2016; O’Brien et al., 2018). Existing cadherins likely mediate weak interactions in low CHIR, which are stabilized leading to cell aggregation on induction in high CHIR. Cadherin switching has been observed in EMTs including the emergence of migratory neural crest (Wheelock et al., 2008) and the progression of cholangiocarcinoma (Araki et al., 2011). A downregulation of Cdh11 has also been observed in differentiation of osteocytes and adipocyte stem cells to more mature cell types (Alimperti and Andreadis, 2015).

Various observational and genetic studies have explored the potential role of cadherin complexes in the morphogenesis of the nephron. Several cadherins have been reported within renal vesicles and later nephron anlagen to the S-shaped body stage including Cdh1 (Cho et al., 1998), Cdh2 (Naiman et al., 2017), Cdh3 (Goto et al., 1998; Lefevre et al., 2017), Cdh4 (Dahl et al., 2002) and Cdh6 (K-cadherin, (Cho et al., 1998). Further, Lefevre *et al*., 2017 catalogued cadherin expression within multiple bulk-mRNA sequencing data highlighting *Cdh2*, *Cdh4* and *Cdh6* in the mesenchymal NPCs in the cap mesenchyme in the developing kidney and *Cdh2*, *Cdh3*, *Cdh4*, *Cdh6*, *Cdh11* and *Cdh16* from renal vesicle to S-shape body stages. These findings are broadly consistent with data here. In addition, the single cell studies underscore the highly dynamic and regional interplay of different cadherins consistent with continuing roles in the nephrogenic program beyond the initial induction and aggregation events that are the focus of this study. Zebrafish studies of spinal cord assembly highlight the important role played by cell type-specific combinatorial expression of different classes of cadherins through create a differential adhesion code in organizing the developing spinal cord in response to a sonic hedgehog morphogen gradient (Tsai et al., 2020).

Despite some considerable efforts to identify developmental roles for cadherins in the kidney either individually or in compound mutant studies (Dahl et al., 2002), the only reported phenotypes have come from antibody interference studies with Cdh6 in kidney explants in which Cdh6 antibodies inhibited the MET of NPCs (Cho et al., 1998). However, analysis of Cdh6^−/−^ mutant kidneys, argued for a different phenotype in renal vesicle polarity and the interconnection of nephrons with the ureteric epithelial network (Mah et al., 2000). Further, an absence of Cdh6 at the cell membrane indicates Cdh6 is not expected to play a role in cell aggregation in the NPC induction model.

Interestingly, altering cadherin levels, in the KO of *Cdh2*, *Cdh4* and *Cdh11*, dramatically alters cell associations within cell aggregates, suggestive of cadherin-directed mechanisms where differing levels, or distinct forms of cadherins, control adhesiveness, cell sorting and self-organization in tissue morphogenesis (Steinberg, 2007). A clustering phenotype has been described in other stem cell systems (Tse et al., 2021). Stem cell-directed human kidney organoids undergo a Wnt-induced MET and subsequent polarization of renal vesicles in response to a brief high CHIR stimulation (Glykofrydis et al., 2021; Morizane and Bonventre, 2017). Thus, the organoid model could be an additional informative system to addressing the question of how a Wnt pulse leads to complex cellular organization (Nishinakamura, 2023) (Nishinakamura, 2019). The organoid model could provide a powerful platform to studying cadherin directed cell association, regional patterning and tissue organization. However, our study highlights the experimental challenge to mechanistic dissection of multiple overlapping cadherin domains.

While loss of β-catenin blocks both induction of the nephrogenic transcriptional program and cell-cell aggregation, loss of Lef/Tcf factors mediating β-catenin’s transcriptional role did not prevent cell-cell aggregation, although Lef/Tcf KO cells adopt a non-random distribution in cell aggregates. Multiple lines of evidence demonstrate that disrupting cell aggregation did not alter the high CHIR and Wnt-mediated transcriptional component of the NPC inductive response. Though α-catenin forms a dynamic linkage between membrane localized cadherin-catenin complexes and the actin cytoskeletal network (Shapiro and Weis, 2009; Yamada et al., 2005), a transcriptional role has been posited for α-catenin in Xenopus development (Sehgal et al., 1997), mammalian cell studies (Daugherty et al., 2014; Giannini et al., 2000). However, our findings that α-catenin-KO NPCs were transcriptionally almost identical to wild-type NPCs argues against a transcriptional role for α-catenin in NPC programs, in line with studies of α-catenin in the brain (Lien et al., 2006). Thus, while β-catenin links the MET to the inductive transcriptional response following Wnt-mediated activation of the nephrogenic program, these distinct cellular processes can be separated genetically and continue independently. A similar concentration of CHIR invoking cell aggregation in NPCs leads to Cdh2 and β-catenin driven aggregation of mouse embryonic stem cells (Sineva and Pospelov, 2010).

Though cadherins undergo heterophilic or homophilic interactions (Takeichi, 2023), our genetic analysis suggests a high degree of functional redundancy amongst cadherins in the aggregation of NPCs. Indeed, the findings underscore the value of the *in vitro* model in enabling the efficient KO of multiple targets, which would be a significant challenge *in vivo*. The similar phenotypes resulting from multiple loss of cadherins and α-catenin points to conventional cadherin complex assembly in the MET, and gene expression studies indicate expression of these cadherin genes is governed by the Wnt/β-catenin driven transcriptional response: Cdh2 and Cdh11 are expressed at substantial levels in naïve NPCs and downregulated on CHIR activation while Cdh3 and Cdh4 are both transcriptionally upregulated on induction. Cdh3 and Cdh4 are targets of the Lef/Tcf/β-catenin transcriptional program (Bugacov *et al*., submitted). Comparison of Cdh2, Cdh4, and Cdh11 KO with KO of all four cadherins argues for a significant role of newly transcribed Cdh3 in the cell adhesion process. Further, *Ctnnd2*, which encodes delta-catenin, a protein bridging and linking cadherin complexes, is also a transcriptional target of Wnt/β-catenin pathway (Bugacov *et al*, submitted; Table 1.1). Thus, the transcriptional program may reinforce cell adhesion and control subsequent morphogenesis of the epithelial nephron.

β-catenin’s dual roles, as a cytoplasmic linker to membrane cadherins and a co-factor in Lef/Tcf transcriptional complexes have been mapped to distinct domains and modifications within the β-catenin protein (van der Wal and van Amerongen, 2020). The dual role of β-catenin raises the question of whether each function is served by unique or common pools of β-catenin (van der Wal and van Amerongen, 2020). Interestingly, in transcriptional profiling following removal of all four cadherins, we do not observe any significant change in the transcriptional response despite the expected freeing of β-catenin associated with membrane cadherin partners. These observations argue against a common pool model. In contrast, the full epithelial transition of aggregated NPCs requires a downregulation of β-catenin transcriptional activity in NPC culture and in *in vivo* nephrogenesis (Park et al., 2012), and non-canonical autocrine signaling by Wnt4, itself a direct target of Lef/Tcf/ β-catenin transcriptional complexes (Guo et al., 2021; Park et al., 2012; Tanigawa et al., 2011);). The block to epithelial formation could be explained by the transcriptional requirement limiting β-catenin availability for the formation of epithelial cadherin complexes.

In *Xenopus* overexpression of cadherins, translocated β-catenin to the cell membrane, inhibiting β-catenin transcriptional activity (Fagotto et al., 1996; Heasman et al., 1994). In *Drosophila* the overexpression of full-length E-cadherin (Cdh2) or its dominant negative truncated form resulted in similar phenotype to the *wingless* (*Drosophila* Wnt-1 homologue) deficient flies (Sanson et al., 1996). Further, in human cell lines, overexpression of soluble Cdh1 and Cdh2 cytoplasmic domains also inhibited transcription of a Lef1 reporter of canonical Wnt transcriptional complexes (Sadot et al., 1998). In these experiments, supra-physiological approaches with abnormally high levels of given factors adds an additional complication to mechanistic interpretation. The *in vitro* NPC model, replicating an *in vivo* event, is well suited to a focused analysis of β-catenin dynamics in cell aggregation and transcription.

Our studies provide a direct and strong linkage of Wnt pathway activation to the MET of NPCs. Interestingly, many studies have highlighted Wnt pathway activation regulating EMTs in normal development and cancer (Arkell et al., 2013; Ji et al., 2019; Schepers and Clevers, 2012; Xu et al., 2020). A potential explanation for these opposing roles lies in the differential targets of β-catenin directed transcriptional programs. In our studies, high CHIR results in *de novo* expression of cadherins Cdh3 which may enhance cell-cell adhesion in the epithelial transition. Importantly, EMT is commonly associated with transcriptional upregulation of the transcriptional repressors Snail/Sna1 and Slug/Sna2, direct targets of canonical Wnt transcriptional complexes in these paradigms (Heuberger and Birchmeier, 2010; Wu et al., 2012; Yook et al., 2006). Snail and Slug are master regulators of transcriptional programs promoting EMT, in part through transcriptional silencing of cadherins stabilizing epithelial organization (Bolós et al., 2003; Cano et al., 2000). Thus, the morphological outcome to canonical Wnt signaling input, MET or EMT, likely reflect distinct, cell type specific, epigenetic programming of Wnt-responsive cells.

## Materials and Methods

### Animals

All animal related research was reviewed and approved by the Institutional Animal Care and Use Committees (IACUC) at the University of Southern California and experimental work been conducted accordingly. Midday on the morning of detection of a vaginal plug was considered to be the E0.5 timepoint. Mouse embryos were recovered for NPCs isolation or histological section analysis at E16.5 stage. WT NPCs were isolated from SWR/J mice. KO experiments have been performed with NPCs derived from crosses of *Gt(ROSA)26Sor^tm1.1(CAG-cas9*,-EGFP)Fezh^*/J (Platt et al., 2014) and SWR/J mice or matings set up with B6.129-*Ctnnb1^tm2Kem^*/KnwJ (Brault et al., 2001). Six2-TGC-tdTomato/Hoxb7-Venus kidneys were generated by crossing male Tg(Six2-EGFP/cre)1Amc/J (Kobayashi et al., 2008) mice with *Rosa26^tdTomato^* (B6.Cg-*Gt(ROSA)26Sor^tm14(CAG-tdTomato)Hze^*/J) females (Madisen et al., 2010). Progeny were crossed with homozygous tgHoxb7-Venus (Tg(Hoxb7-Venus*)17Cos/J) females (Chi et al., 2009). For generating Vid. S3 (KO experiment with high resolution imaging of membrane fluorescent reporter) mice constitutively expressing Cas9-eGFP strain (Platt et al., 2014; IMSR_JAX:026179) were crossed with mTmG fluorescent reporter mice (Muzumbar et al., 2007; *t(ROSA)26Sor^tm4(ACTB-tdTomato,-EGFP)Luo^*/J).

### NPC isolation and culture

NPEM formulation and NPC isolation protocols were based on protocols of (Brown et al., 2015), with modifications by Bugacov *et al*., (submitted). After completing NPC isolation, NPCs were plated in 24 well CELLSTAR® cell culture plates (for bulk-RNA-seq. and live imaging; VWR, 82050-892), µ-Slide 8 Well (for Video S1.; Ibidi, 80827) or on treated 24 well µ-plates (for IF staining; Ibidi, 82426). Plates were coated with Matrigel (Corning, 354277) as described in Bugacov *et al*., submitted.

Seeding densities:

- 16-24 h stabilization, induction for 24 h: 300 000 cells/well 16-24 h stabilization, 24 H KO, induction for 24 h: 100 000 – 150 000 cells/well
- 16-24 h stabilization, 48 h KO: 150 000 cells/well
- 16-24 h stabilization, 48 h KO, induction for 24 h: 50 000 – 100 000 cells/well
- Timelapse in µ-Slide 8 well chip: 16-24 h stabilization, induction for 24 h: 100 000 cells/well
- Timelapse in 24 well plate: 16-24 h stabilization, 24 h KO, induction for 24 h: 150 000 cells/well

For the extracellular Ca^2+^ removal experiments, the following types of cell culture media have been used with supplemented low (1.25 μM) or high (5 μM) CHIR concentrations: DMEM/F-12, HEPES (Thermo Fisher Scientific, 11330-032), DMEM, high glucose, no glutamine, no calcium (Thermo Fisher Scientific, 21068028).

Wnt3a was manufactured by R&D Systems (1324-WN-010/CF) and used at 200 ng/ml for 24 h induction. Bispecific antibody (the gift of the Garcia laboratory; (Janda et al., 2017)) was added at the indicated concentration.

### *In vitro* mRNA synthesis and cell transfection

Cre and mCherry mRNA synthesis and NPC lipofection followed the protocol described in detail by Bugacov *et al*., (submitted). In this set of experiments, we added 500 ng of total mRNA (per transcript type) to 1 well of a 24 well plate in conjunction with sgRNA(s) in the KO experiments. For Vid. S3 NPCs were transfected with Cre mRNA.

### CRISPR mediated gene removal

For KO experiment protocols see Bugacov *et al*., submitted. sgRNAs utilized for 48 h KO period experiments were ordered from Synthego designed by CRISPR Design Tool (except GFP and *Ctnnb1*). Top four guides were triaged based on IF staining KO efficiency. Synthesized gRNAs were reconstituted at 100 pmol/μl in TE buffer and stored at −20 °C. Total sgRNA concentration for the KO experiments was 7.5 pmol/well for a 24-well plate. For the QCKO experiments, 1. 875 pmol of each gRNA was added to each well and 2.5 pmol of each gRNA for triple KO Cadherin experiments (*Cdh2/4/11*). mCherry mRNA was co-transfected at a concentration of 500 ng/well. All sgRNA sequences are listed in **Table 2.1**.

### FACS sorting, mRNA isolation and RT-qPCR

For detailed protocols see Bugacov *et. al*, submitted.

### Bulk RNA-sequencing

Library preparation and methods for data analysis follow procedures in Bugacov *et al*., (submitted). To identify differentially expressed gene lists, normalized counts tables were inputted to DeSEQ2 (Love et al., 2014) using a Log_2_FC cut off +1 or −1, and an adjusted p-value no greater than 0.05. ggplot and complex heatmap functions in R (Gu et al., 2016) were used to visualize data and Benjamini-Hochberg correction (False Discovery Rate) was implemented with gene normalized counts greater than or equal to 10 using a hypergeometric test.

### Immunofluorescent analysis

Immunofluorescent analysis of proteins followed the procedure outlined by Bugacov *et al*., submitted with modifications. NPCs were cultured on tissue culture treated 24 well µ-plates (Ibidi, 82426) and washed with pre-warmed PBS and fixed with ice-cold 4% PFA in PBS on ice for 30 min, then washed with PBS twice. For time series experiment, fixation happened at 0, 6, 12 and 24 h timepoints. For nuclear labeling, cells were incubated with 1:10000 Hoechst 33342 (Thermo Fisher Scientific, H3570) in PBS for 10 mins and finally solution was changed to PBS. 24 well plates were kept away from light at 4 °C prior to image acquisition. Primary and secondary antibodies are listed in **Table 2.2**.

### Image acquisition and quantification

Phase contrast images were acquired by DLPlan Fluor 10x/NA 0.3 dry, Ph1 DL objective connected to Mono Camera Nikon DS-Fi3 in Nikon ECLIPSE Ts2R inverted research microscope (Nikon Instruments). Confocal images were collected using Leica SP8-X confocal fluorescence imaging system (Leica Microsystems) with 40x/1.3 NA oil, HC PL APO CS2 or 63x/NA 1.4 oil, HC PL APO CS2 objectives in 1024−1024 pixels. Brightfield images was acquired by Leica Thunder widefield microscope (Leica Microsystems) with 40x/NA 0.60 dry, HC PL FLUOTAR L objective. When area of interest did not fit into one scanning plane, Z-stacks were acquired and represented as stacked images.

Timelapse recording for Vid.S1 was recorded with 40x/1.3 NA oil, HC PL APO CS2 on Leica SP8-X, Vid.S2. and S3 was acquired by 10x/NA 0.45 dry, HC PL APO or 40x/NA 0.60 dry, HC PL FLUOTAR L on Leica Thunder widefield microscope, respectively. During timelapse image acquisition homeostatic conditions were maintained with Ibidi stage top incubation system (Ibidi, 10720).

Images were quantified (“surface”, “cell”, “spots” modules) in Imaris microscopic image analysis software (version 10.0, Oxford Instruments). Procedures applied to quantifying the removal of membrane α-catenin, Cdh2, Cdh3, Cdh4, and Cdh11 proteins are detailed in Bugacov *et al*., submitted. Fluorescent intensity histograms were created for “surface”, “cell” and “spots” modules for mCherry channel to classify mCherry^+^ and mCherry^−^ NPCs based on steep intensity drop offs in the histogram. The criteria for assessing Jag1^+^ cells is detailed in Bugacov *et al*., submitted.

“Spots” module was used to track the NPCs during Vid.S1., to determine the number of mCherry^+^ cells outside the aggregates, and to measure distance to nearest neighbor (cells were excluded from evaluation if their nearest neighbor were more distant than mean + 2SD at Fig1. F.). Cell tracks in Vid.S1., cell-cell contact numbers and durations were manually annotated, as well as membrane and filopodia contact events (see examples on Fig1. SE). Cellular height, cellular volume, cellular size and nuclear height, nuclear volume were quantified by “surface” or “cell” modules.

Cells were considered sorted from cell aggregates if the cell body showed no membrane or filopodial contact with the Jag1^+^ NPC aggregate. Clusters within the aggregates of Cdh2+4+11 KO experiments were defined based on the following criteria: 1) at least five transfected cells were adjacent to each other; 2) a maximum of one non-transfected cell with the transfected cell cluster; 3) the cluster was entirely surrounded by non-transfected cells and/or the cluster was positioned at the boundary of an aggregate. Clusters were manually annotated and counted with the “spots” module.

We applied median filter and background subtraction in Imaris for Vid.S1. Vid.S2. and S3. were processed by computational clearing, median filtering and thresholding in LASX. The frames of Vid.S3. have been stabilized in ImageJ.

### Statistics and data plotting

We evaluated the normal distribution of datasets by D’Agostino-Pearson test. When we compared two independent groups with a normal distribution, significance levels were determined by an unpaired *t* test. With a non-normal distribution, we utilized a Mann-Whitney test. Comparing more than two groups with a normal distribution, we performed a one-way ANOVA. With multiple samples and a non-normal distribution, p-values were calculated with a Kruskall-Wallis test. Mixed effect statistical analysis was applied for Fig. S1D and 2H datasets. Data is plotted as mean ± SD unless otherwise specified in figure legends. Statistical tests were considered significant with p of 0.05 or less. Graphs were plotted by Prism 9.4 (GraphPad by Dotmatics). Summary schematics were created with BioRender.com.

## Supporting information

Vid. S1

Vid. S2

Vid. S3

## Acknowledgments

We are grateful to Dr. Seth Ruffins in the Eli and Edythe Broad CIRM Center Optical Imaging Facility for his help with image acquisition and microscope setup. In the McMahon lab, we thank for Jin-Jin Guo for assistance with mouse husbandry and technical help with sgRNA testing, Sunghyun Kim and Cheng Jack Song for helpful discussions, and Muskaan Singh for data collection. We thank Dr. Christopher Garcia and Dr. Yi Miao for sharing the Wnt mimetic bi-specific antibody.

## Author contributions

BD, HB and APM conceived the study. BD, HB and B-MB performed experiments and collected data, BD, HB, and B-MB analyzed the data and BD, HB and APM wrote the manuscript.

## Diversity and inclusion statement

In order to advance our educational, scientific and clinical mission, USC Stem Cell is deeply committed to creating a culture that supports diversity, equity and inclusion (DEI). HB identifies as an underrepresented minority female in STEM and benefitted from funds from the Ruth L. Kirschstein National Research Service Award (NRSA) Individual Predoctoral Fellowship to Promote Diversity in Health-Related Research (Parent F31-Diversity) and F31 DK122777.

---

**Figure S1.**
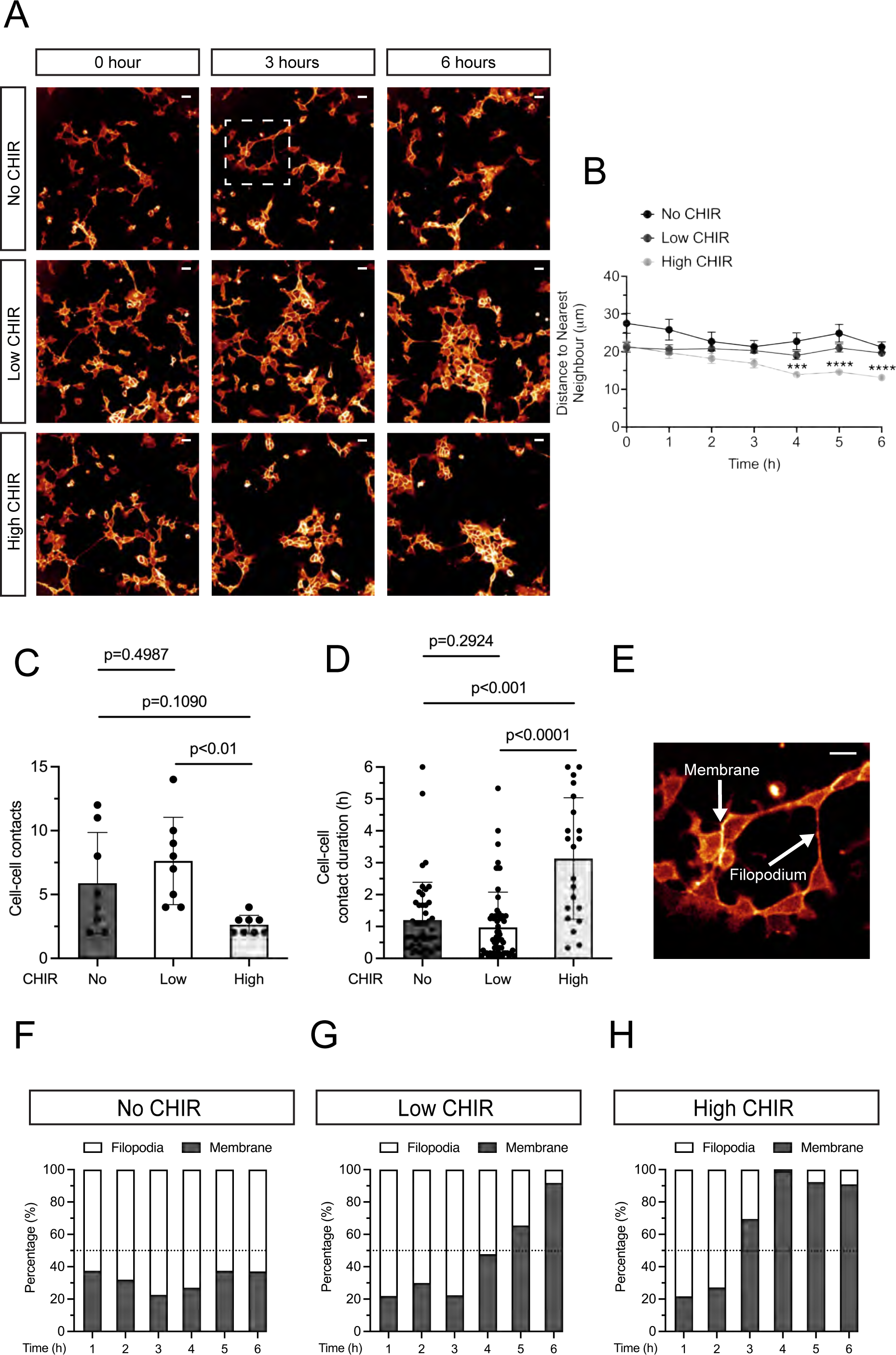
Detailed analysis of individual NPC behavior with increased Wnt stimulus. (A) Time lapse stills of NPCs with membrane tdTomato fluorescent reporter cultured in no (0 μM) CHIR, low (1.25 μM) CHIR and high (5 μM) CHIR over 6 h. Images are derived from Vid.S1. Squared area is magnified in Fig. S1E. Scalebars are 20 μm. (B) Quantification of distance to nearest neighbor cell over the culture period above. Statistical analysis: mixed effect analysis. Graph is plotted as mean ± SEM. For each field of view or condition cell position was annotated manually. Average evaluated number of cells/field of view: no CHIR=49, low CHIR=93, high CHIR=84). (C–D) Quantification of contact numbers per cell and contact duration in NPC cultures. Datapoints represent eight individual cells at starting timepoint field of view/condition (C), the intercellular contact durations of these eight cells are displayed in (D) Statistical tests were ordinary one-way ANOVA and Kruskall-Wallis test, respectively. (E) Representative images of the types of cell-cell contacts: filopodium or membrane-membrane contacts. (F–H) Quantification of filopodia and membrane-membrane contact ratio over 6 h in no CHIR, low CHIR and high CHIR conditions. The dotted grid line labels 50%.

**Figure S2.**
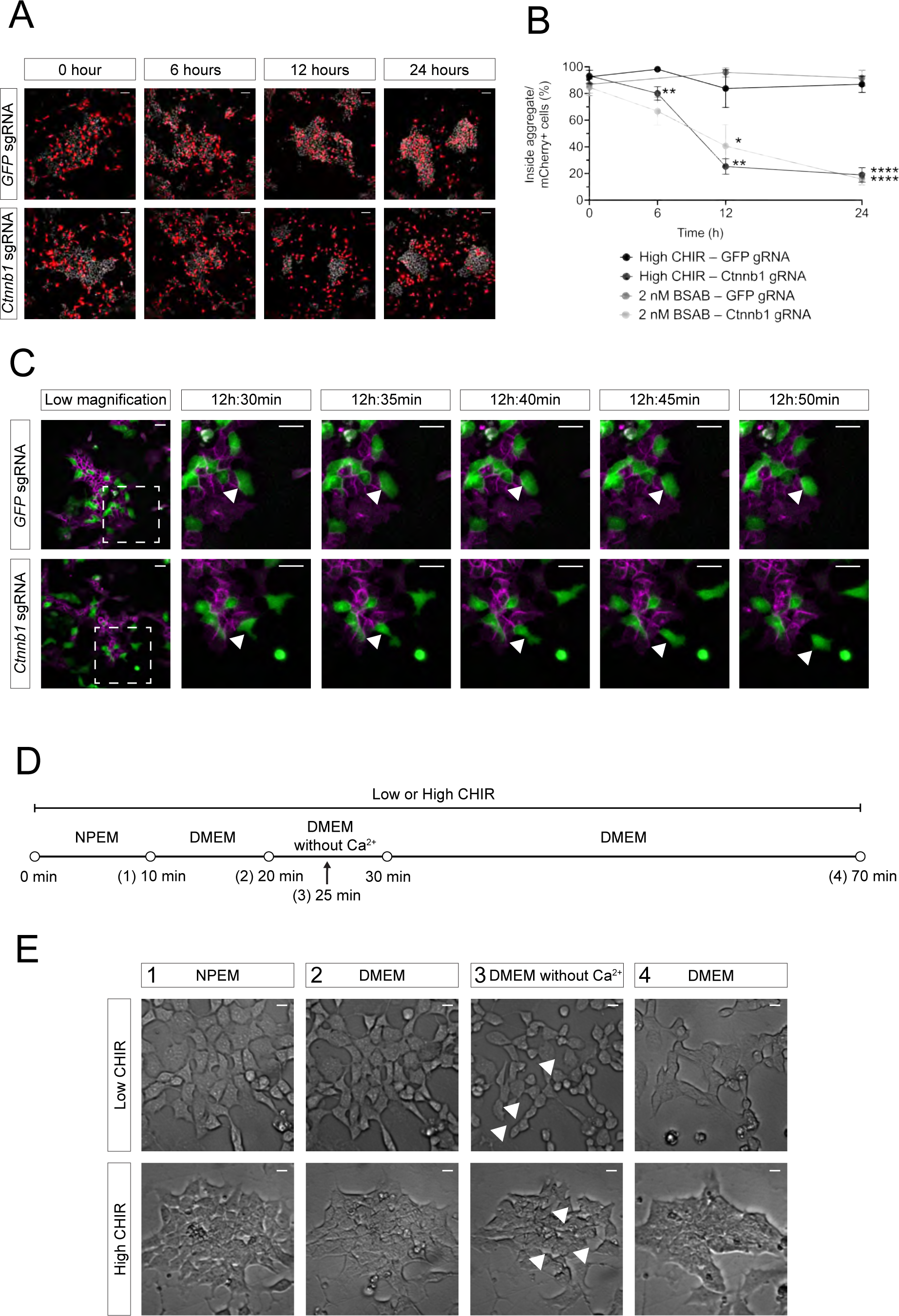
Detailed analysis of cell sorting following ß-catenin KO and Response of NPCs to Ca^2+^ removal. (A) Representative images of time series experiment of CTRL (GFP sgRNA) and β-catenin-KO (Ctnnb1 sgRNA) conditions when NPCs were fixed and IF stained at 0, 6, 12 and 24 h after the initiation of induction. Scale bars are 50 μm. (B) Quantification of time series experiment by % of cells within aggregates. n=5–10 field of views/well, 1–2 biological replicates, 1–2 technical replicates, mixed effect statistical analysis. (C) Still images showing individual cell behaviour during aggregation and sorting from timelapse Vid. S3. NPCs isolated from mTmG x Cas9-eGFP mice are co-transfected with Cre mRNA and sgRNA targeting GFP (CTRL) and β-catenin (Ctnnb1). Scale bars are 10 μm. (D) Schematic representation of the experimental protocol to examine the role of extracellular Ca^2+^ in NPC culture. (E) Representative images of E16.5 NPCs cultured in NPEM (CTRL), DMEM (negative CTRL), DMEM without Ca^2+^ (experimental) and the re-addition of DMEM in low and high CHIR conditions. The loss of cell-cell contacts are labelled with white arrowheads. Scale bars are 10 μm.

**Figure S3.**
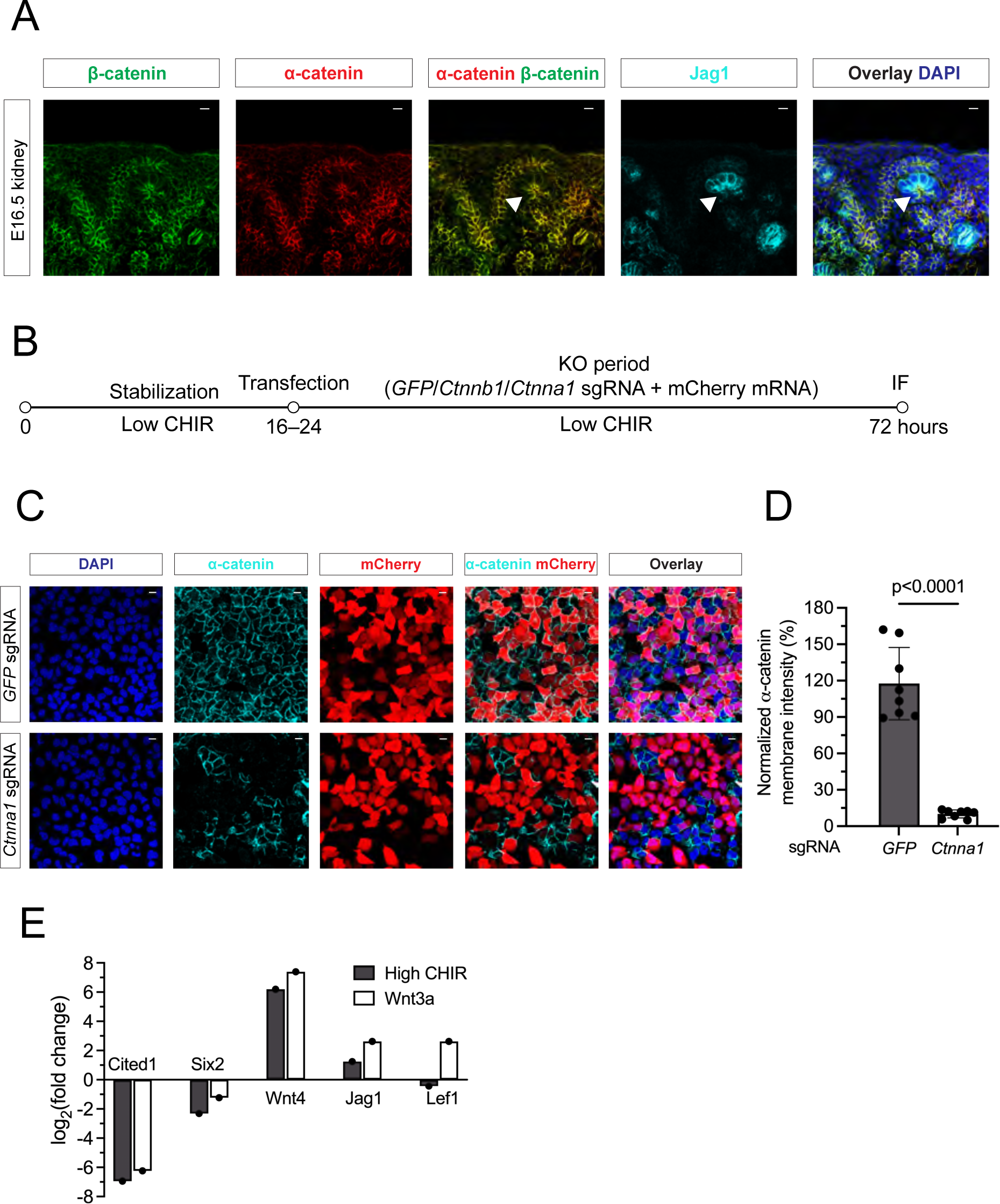
In vivo characterization of α-catenin and in vitro validation of sgRNA-Cas9 system for CTRL and α-catenin-KO condition. (A) Representative images of IF co-staining of α- and β-catenin (α-catenin: green, β-catenin: red) and the induction marker Jag1 (cyan) in wild-type E16.5 kidney. Arrows mark RV with apical accumulation of catenins and strong distal expression of Jag1. Scale bars are 10 μm. (B) Schematic representation of the experimental protocol to validate removal of α-catenin. (C) Representative IF images of α-catenin removal (cyan) in mCherry transfected cells (red) with nuclear staining DAPI (blue). Scale bars are 10 μm. (D) Quantifications of the membrane intensity of α-catenin KO cells. Unpaired *t* test. (E) RT-qPCR dataset showing comparable upregulation of induction genes and the downregulation of self-renewal gene network after 24 h of incubation with high CHIR and Wnt3a (200 ng/ml).

**Figure S4.**
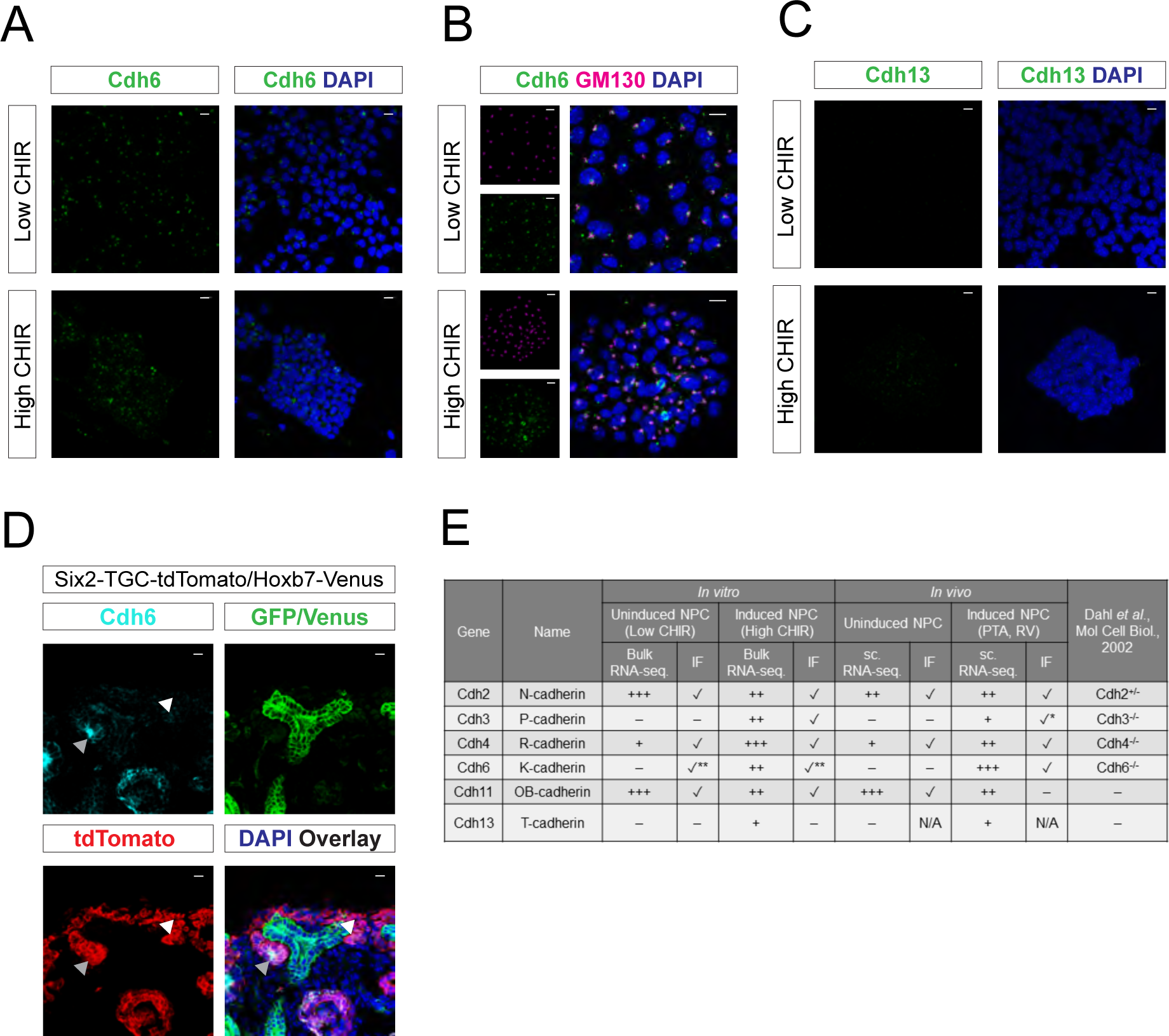
In vitro and in vivo analysis of mRNA and protein levels for additional cadherins. (A) IF staining of Cdh6 (green) in isolated E16.5 NPCs shows a punctate pattern of protein localization in low and high CHIR conditions. (B) Co-labeling of Cdh6 and GM130 Golgi-marker (Cdh6: green, GM130: magenta) in NPCs from E16.5 kidneys cultured in low and high CHIR conditions. Scale bars are 10 μm. (C) IF staining of Cdh13 (green) in isolated E16.5 NPCs shows weak, sporadic membrane localization restricted to high CHIR conditions. (D) Representative images of IF staining of E16.5 Six2-TGC-tdTomato/Hoxb7-Venus mouse kidneys highlighting indicated cadherins (cyan), tdTomato (nephron lineage, red) and GFP: nuclear GFP highlights Six2 in NPCs while membrane GFP labels Venus reporter in the ureteric lineage. Cdh6 is not detected in uninduced NPCs but Cdh6 is present in the late RV (white and gray arrowheads). (E) Table summarizing *in vitro* and *in vivo* cadherin expression and protein levels in E16.5 mouse NPC culture and the E16.5 mouse kidney. (–) no expression/presence of mRNA/protein. + to +++: low to high expression/levels of mRNA or protein. ✓*: protein is present in distal RV. ✓ **: protein is presented in Golgi-apparatus. RV: renal vesicle, PTA: pre-tubular aggregate

**Figure S5.**
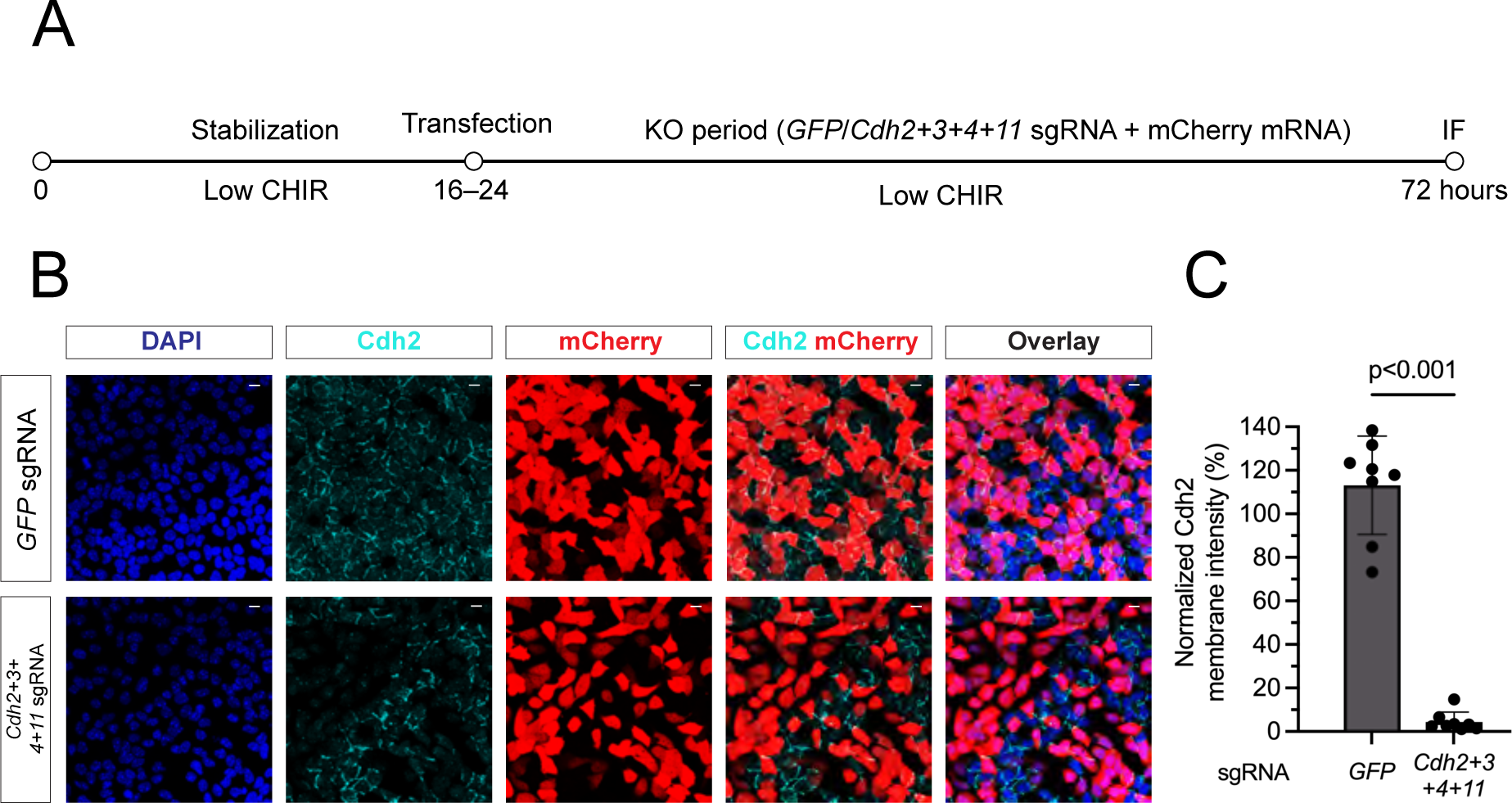
Confirming the removal of Cdh2 protein removal in QCKO by the Cas9-sgRNA system. (A) Schematic representation of the experimental protocol to remove Cdh2+3+4+11 including 48 h KO period. (B) Representative IF images of Cdh2 removal (cyan) in mCherry transfected cells (red) with nuclear staining DAPI (blue). Scale bars are 10 μm. (C) Quantifications of the membrane intensity of Cdh2 in QCKO cells. Mann-Whitney test.

**Figure S6.**
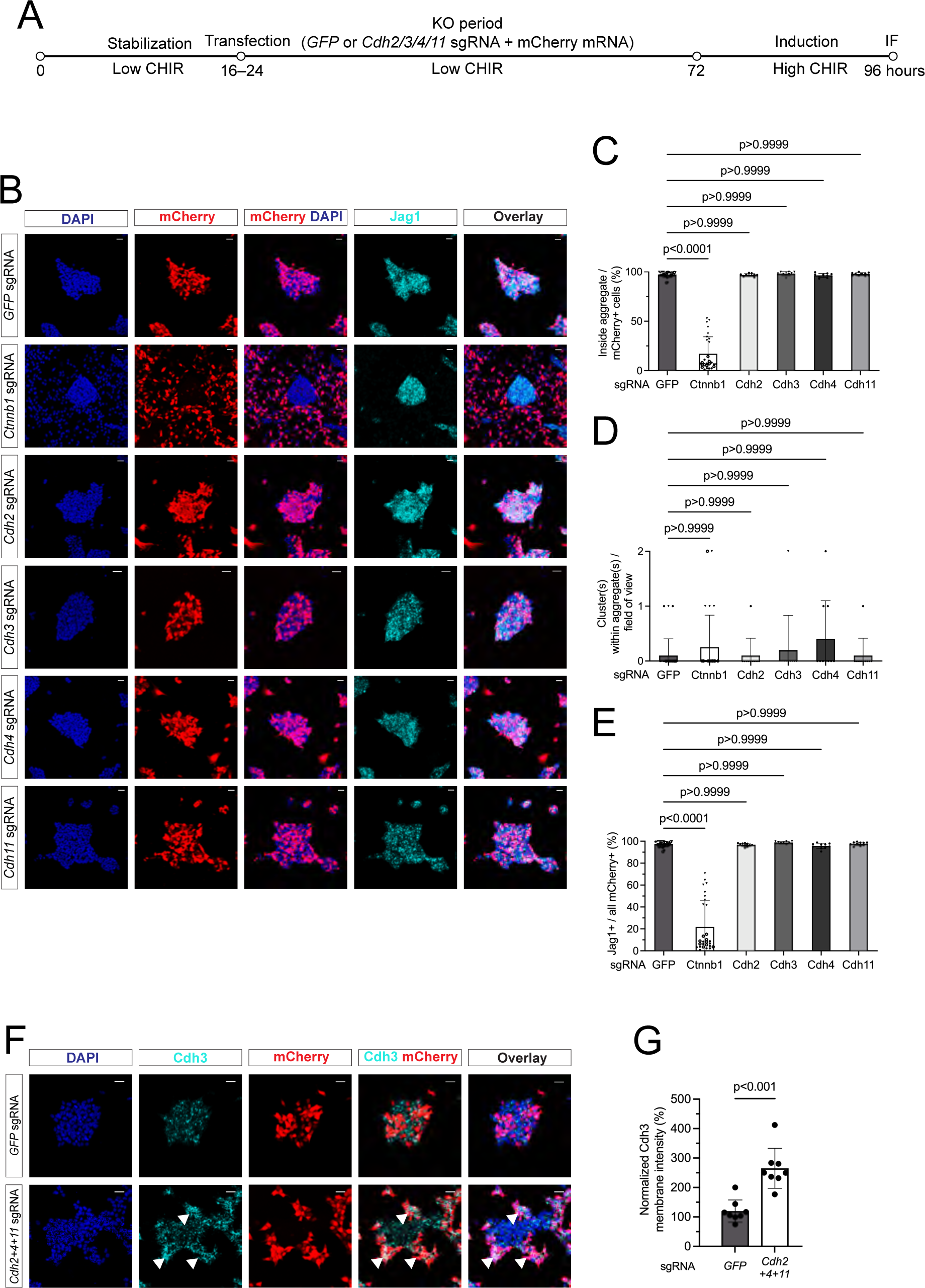
Individual cadherin removal does not influence high CHIR-dependent cell aggregation or induction of transcriptional targets. (A) Schematic of the experimental protocol to investigate the effects of individual removal of Cdh2, Cdh3, Cdh4 or Cdh11 from E16.5 NPC cultures in high CHIR conditions. (B) Representative images of negative CTRL GFP sgRNA, positive CTRL β-catenin-KO and individual cadherin KO in E16.5 NPC culture in high CHIR: DAPI, nuclei; mCherry^+^, transduced NPCs, Jag1^+^, induced NPCs. Scale bars are 25 μm. (C–D) Quantification of changes in cell clustering of transduced NPCs. (C) percentage of transfected cells within aggregates. (D) number of within cell aggregate clusters of transfected NPCs. Biological replicates are represented by different symbol shapes and technical replicates are highlighted by different fill colors. Statistical analysis using a Kruskall-Wallis test. (E) Percentage of Jag1^+^ transfected NPCs in cadherin KOs. Biological replicates are represented by different symbol shapes and technical replicates are highlighted by different fill colors. Statistical analysis using a Kruskall-Wallis test. (F) Representative images of Cdh3 IF staining GFP sgRNA and β-catenin-KO showing elevated membrane intensity of Cdh3 (cyan) on Cdh2, Cdh4 and Cdh11. Scale bars are 25 μm. (G) Quantification of the membrane intensity of Cdh3 in CTRL and Cdh2, Cdh4 and Cdh11 KO cells. Mann-Whitney test.

## Supplementary videos

Video S1. Timelapse recordings showing individual NPC behavior in no (0 μM), low (1.25 μM) and high (5 μM CHIR) conditions: Fig. S1 shows snapshots from this video data. NPCs are isolated from mT/mG mice and show membrane tdTomato reporter activity. Scale bars are 10 μm.

Video S2. Timelapse recordings showing NPC cell sorting comparing mCherry transfected NPCs (red) receiving either GFP sgRNA (CTRL) or *Ctnnb1* sgRNA (β-catenin-KO). mCherry signal is projected on the brightfield channel. Recording covers 24 h period following addition of high CHIR. Scalebars are 100 μm.

Video S3. Timelapse recordings showing high-resolution views of cell sorting by β-catenin-KO cells in high CHIR (views related to Figure S2C). Non-transfected cells are magenta and transfected NPCs are green. Arrows highlight representative cellular events. Scalebars are 10 μm.

## Supplementary tables

Table S1. Summary table of bulk RNA-sequencing data corresponding to Fig. 7.

Table S2. Details of sgRNA manufacturers and sequences, and primary and secondary antibodies.

